# Mesenchymal stem cell-derived extracellular vesicles alleviate imiquimod-induced psoriasis symptoms in a mouse disease model

**DOI:** 10.1101/2024.09.10.612372

**Authors:** Yuan Yi, Yu Yan, Guowu Liu, Chuang Cui, Yanhua Zhai, Xinjun He, Ke Xu

## Abstract

Psoriasis is an immune mediated, chronic inflammatory skin disease. Mesenchymal stem cell-derived extracellular vesicles (MSC-EVs) have inherent immunomodulatory potency from their parental cells, the mesenchymal stem cells (MSCs). In this study, we revealed the application potential of MSC-EVs to alleviate psoriatic symptoms in imiquimod-induced psoriasis mouse model.

## Introduction

Mesenchymal stem cells (MSCs) are pluripotent stem cells with the ability to self-renewal and multi-potential differentiation. MSCs are found in a variety of tissues such as bone marrow, adipose tissue, umbilical cord blood and placenta^1^. MSCs function through a variety of mechanisms. First of all, mesenchymal stem cells have the potential for multi-directional differentiation and can be differentiated into many different types of cells, such as osteocytes, chondrocytes, muscle cells and adipocytes, with a wide range of applications in tissue engineering and regenerative medicine^2^. In addition, MSCs secret bioactive factors (cytokines, growth factors, chemokines, pro-angiogenesis and antioxidants) to reduce the stress response, modulate apoptosis and promote tissue repair and regeneration^3^. MSCs also play a role in regulating local and systemic inflammation, which can inhibit excessive immune responses and reduce inflammation^4^. These mechanisms have led to the potential applications of MSCs in a wide range of disease areas, such as graft-versus-host disease, Alzheimer’s disease, diabetes, cardiovascular diseases and many other diseases^5-7^.

Mesenchymal stem cell-derived extracellular vesicles (MSC-EVs) are nanoscale membrane vesicles secreted by MSCs. These vesicles are typically between 30-150 nm in diameter and have a bilayer phospholipid membrane structure that contains a variety of bioactive substances such as proteins, mRNAs, miRNAs and lipids, playing important roles in cell-to-cell communication and regulation of physiological functions^8^. MSC-EVs have the property of low immunogenicity, targeting ability, anti-fibrotic and anti-apoptotic effects and immunomodulatory function, empowering MSC-EVs a wide range of potentials in the treatment of different diseases^9^. MSC-Evs can reduce cardiac fibrosis, improve cardiac function, promote cardiomyocyte regeneration, therefore they have great value in the treatment of cardiovascular diseases such as myocardial infarction and heart failure^10^. MSC-EVs exhibit immunomodulatory effects in the treatment of autoimmune diseases such as rheumatoid arthritis (RA)^11^, multiple sclerosis (MS)^12^ and systemic lupus erythematosus (SLE)^13^. In addition, MSC-EVs can reduce inflammation and promote lung tissue repair, and have therapeutic effects in respiratory diseases such as chronic obstructive pulmonary disease (COPD), asthma, and pulmonary fibrosis^14^. In neurological diseases such as stroke, spinal cord injury, and Alzheimer’s disease, MSC-EVs contribute to neuroprotection and regeneration^15^. In fracture healing and cartilage injury repair, MSC-EVs help promote the regeneration of bone and chondrocytes^16^. Previous researches have shown that MSC-EVs can inhibit tumor growth and metastasis as well^17^.

## Results

### MSC-EVs ameliorated the symptoms in a mouse model of imiquimod (IMQ) induced psoriasis

EVs secreted from MSCs have been shown to have therapeutic effect in inflammatory diseases. In this study, we topically applied MSC-EVs in an IMQ induced psoriasis mouse model. First of all, we isolated EVs from MSC microsphere carrier culture supernatant and monitored particle size and concentration by nanoparticle tracking analysis (NTA), then assessed the morphology of EVs by transmission electron microscopy (TEM) (**Figure 1**). We found that MSC-EVs showed the typical characteristics of cell EVs both in size distribution and TEM imaging.

**Figure 1.**
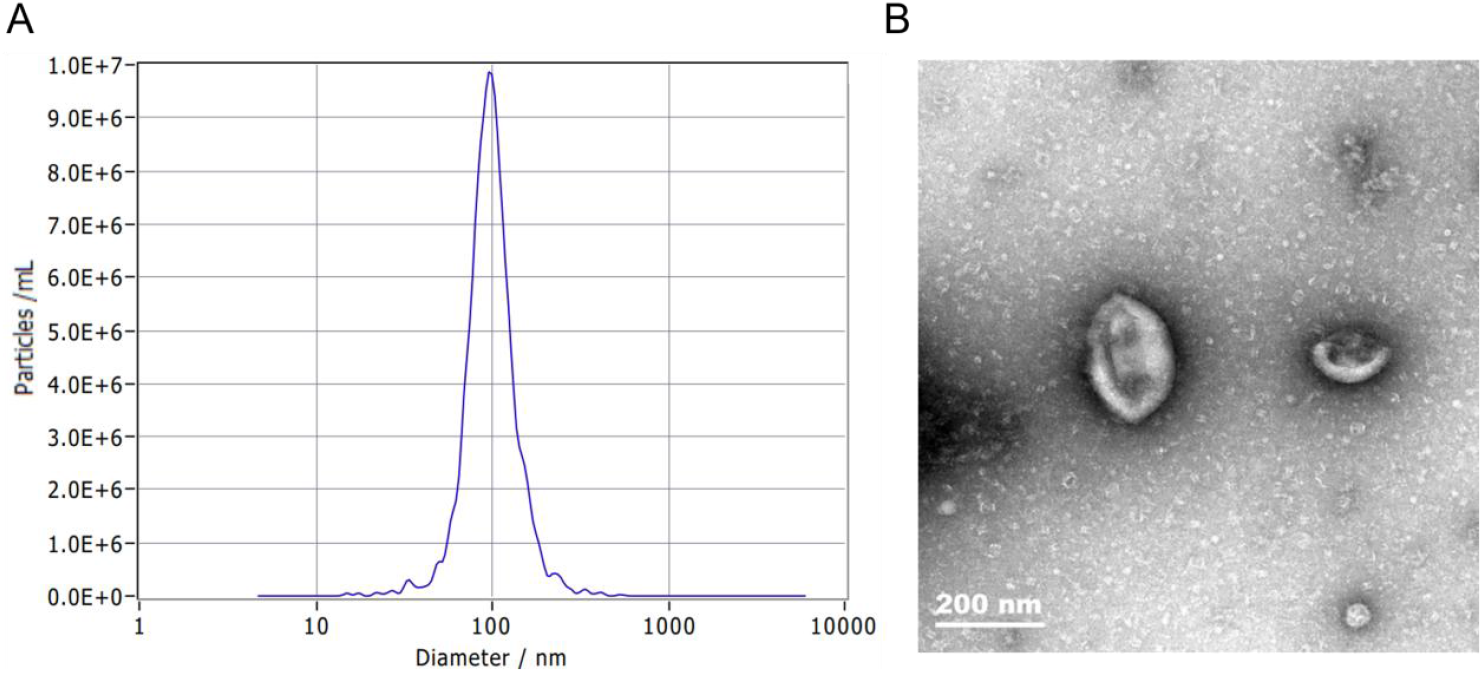
Preparation and analysis of MSC-EVs. (A) The MSC-EVs size distribution and concentration map on NTA. (B) TEM images of MSC-EVs, size bar=200 nm.

To test the therapeutic effects of topically applied MSC-EVs on psoriatic skin, we constructed a mouse model of IMQ induced psoriasis. We applied IMQ to mouse dorsal skin and ear from day 0 to day 5, together with the application of MSC-EVs on the right side dorsal skin and right ear after each IMQ treatment for testing. At the termination (day 5), skin flakes were observed on the left skin and left ear with IMQ treatment, whereas MSC-EVs application could reduce skin flakes of right side skin and right ear (**Figure 2**).

**Figure 2.**
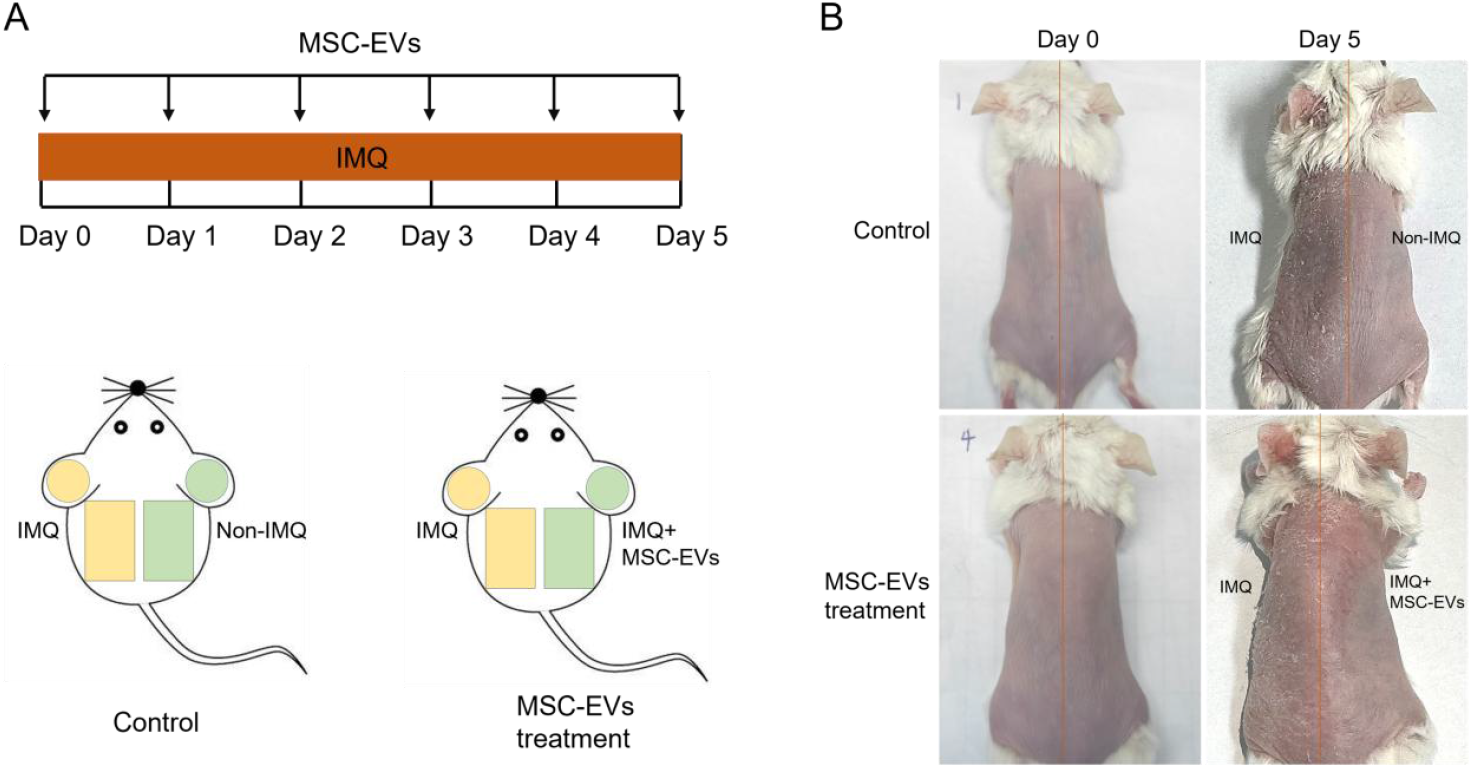
Schematic illustration of the experimental design. (A) Mice were randomized into two groups. In the control group, the left dorsal skin and left ear of each mouse were applied with IMQ cream from day 0 to day 5, while the right dorsal skin and right ear were not treated. In MSC-EVs testing group, both sides of the dorsal skin and ears were applied with IMQ cream from day 0 to day 5, after IMQ was completely absorbed and dried, MSC-EVs were applied into the right dorsal skin and ear. (B) Photographs of mice at day 0 and day 5 before sample collection.

Skin and ear tissues from different treatment were analyzed to investigate the pathological change. In non-IMQ treated control mice, right side dorsal skin and right ear had unimpaired structure, while IMQ application resulted in nucleated stratum corneum and thickened epidermis and dermis. Whereas MSC-EVs was highly effective in restoring the histopathologic changes as indicated with the skin thickness (**Figure 3**).

**Figure 3.**
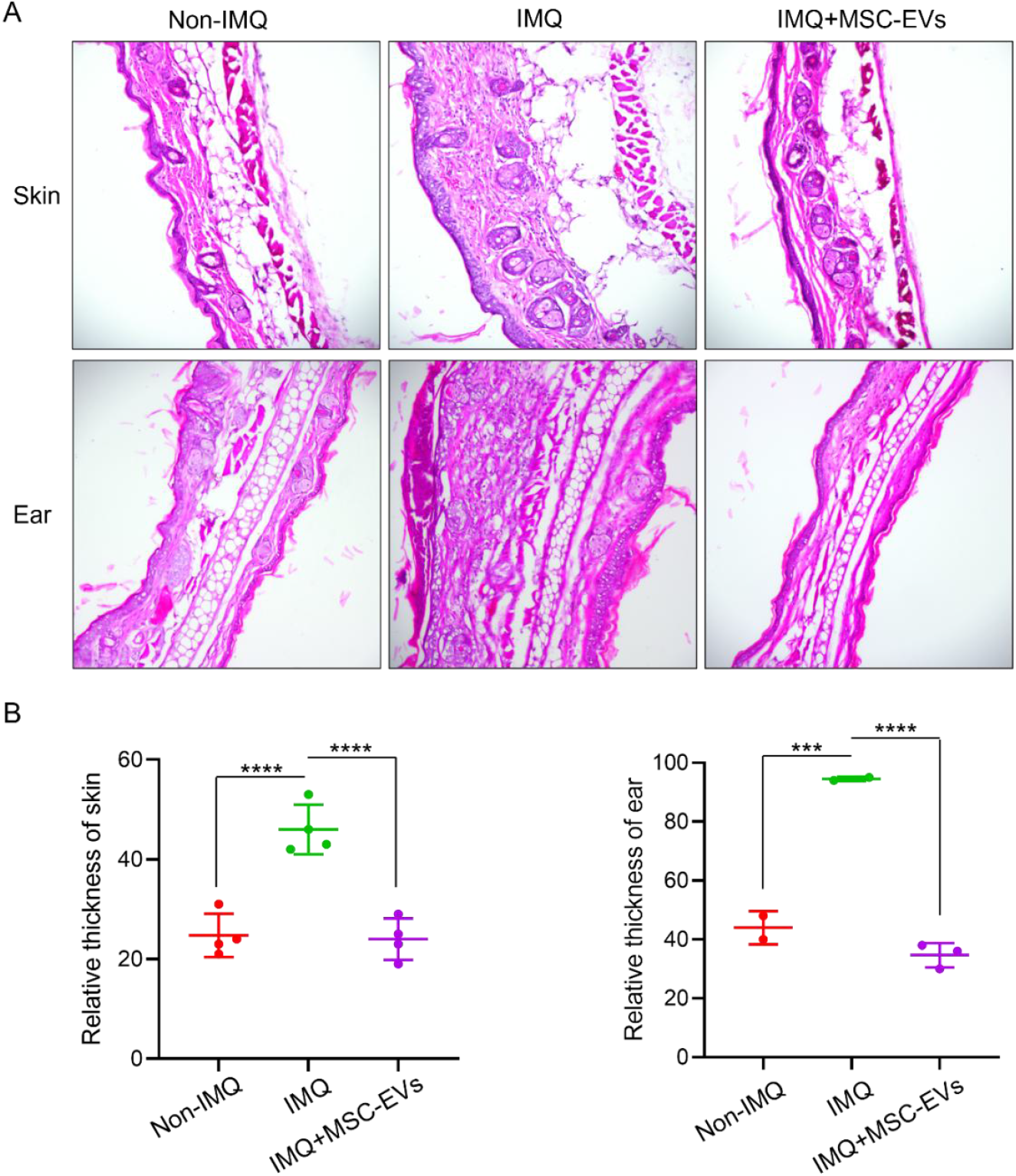
H&E-stained skin and ear sections showed topically applied MSC-EVs alleviated psoriatic skin thickness. (A) Non-IMQ treated mice showed normal structures, IMQ-applied mice had nucleated stratum corneum and widen thickness of epidermis and dermis. whereas MSC-EVs significantly restored skin thickness and maintained skin normal morphology with IMQ-induced psoriasis. Images were taken at 20× magnification. (B) Quantification of skin and ear showing increased thickness in IMQ-applied side compared to normal side, and IMQ induced side with MSC-EVs treatment. Data were presented as mean± SEM. \*\*\**P* < 0.001, \*\*\*\**P* < 0.0001. One-way analysis of variance (ANOVA) was used for the comparisons through using GraphPad Prism 8 software.

Psoriasis is an inflammatory disease, immune cells and cytokines have been involved in pathogenesis of psoriasis. To investigate whether macrophages are implicated in psoriatic immune pathology, we future explored the expression of macrophage marker F4/80 in skin and ear sections. IMQ application induced an increased expression of macrophage marker and promoted their infiltration into dermis. MSC-EVs restored macrophage infiltration and near-normal distribution as shown in IMQ applied psoriasis side compared with the side of non-IMQ application (**Figure 4**).

**Figure 4.**
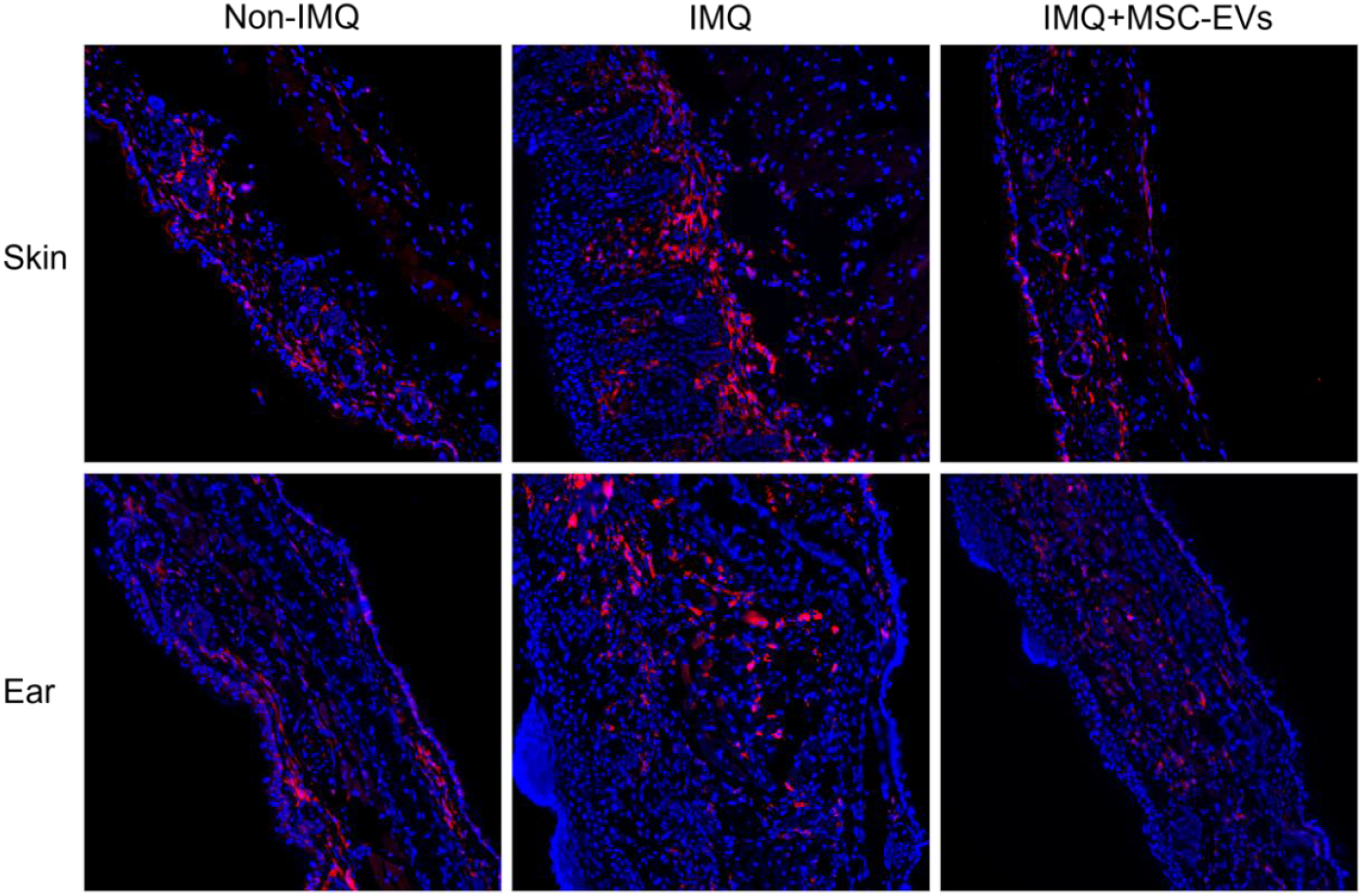
MSC-EVs alleviated macrophage infiltration in dermis. Immunofluorescence labeling of macrophage marker F4/80 showed macrophages were increased and mainly distributed in dermis with IMQ-applied skin, whereas macrophages in mice with MSC-EVs treatment were widely distributed in epidermis and dermis. Images were taken at 20× magnification.

## Discussion

In this study, MSC-EVs have shown their capacity to alleviate symptoms in IMQ-induced psoriasis. We demonstrated that topically applied MSC-EVs could restore skin integrity, thickness and macrophage distribution.

There are still remaining questions to be answered as what the underlying molecular mechanisms are involved in positive effect of MSC-EVs in this animal model. It will be critical to explore the major psoriatic cytokines, such as IL-17 and IL-23 to understand how these cytokines are involved in the treatment of psoriasis. Moreover, as topically applied MSC-EVs is considered inefficient to overcome the skin barrier, it would be interesting to study if the assistance of appropriate administration strategies such as microneedle or subcutaneous injection can effectively increase the efficacy.

## Materials and Methods

### Isolation and identification of MSC-EVs

At the end of cell culture, the culture supernatant was collected and centrifuged at 120 g at 4 °C for 5 min to remove cell precipitation, the clarified supernatant was filtered through 0.22 μm filter membrane (Millipore) to remove cell debris and large vesicles, then the supernatant was ultracentrifuged for 60 min at 133,900 g at 4 °C to isolate MSC-EVs, the EV pellet was finally resuspended in PBS.

The morphology of EVs was observed by transmission electron microscopy (TEM) (FEI) according to the manufacturer’s protocol. EV solution was diluted in PBS to a final volume of 1 mL, size distributions and concentrations were measured by nanoparticle tracking analysis (NTA) on a ZetaView (ParticleMetrix), all settings were set according to the manufacturer’s software manual.

### Imiquimod induced psoriasis mouse model

Female, 8-weeks-old Balb/c mice were purchased from GemPharmatech Co., Ltd. On day 0, the dorsal skin was shaved with depilatory before application of IMQ cream. A psoriatic phenotype was induced by topical application of IMQ on the dorsal skin and ear daily, the right dorsal skin and right ear were then treated with MSC-EVs after the application of IMQ. On day 5, the dorsal skin and ear were collected from each mouse and fixed in 4% paraformaldehyde (PFA) immediately. Non-IMQ induced skin and ear without MSC-EVs treatment as negative control, only IMQ applied skin and ear were collected as disease model controls.

### Histological staining

Skin and ear were fixed in 4% PFA overnight, dehydrated in 30% sucrose solution for 3 days, embedded with cryoprotective medium OCT (SAKURA) and sectioned for 5 μm thickness. Histopathological staining was performed with Hematoxylin & Eosin (H&E) staining kit (Beyotime) according to manufacturer’s protocol. For immunofluorescence staining, primary antibodies against F4/80 (Abcam) were incubated overnight at 4°C and the secondary antibodies (Invitrogen) were incubated for 1 h at room temperature. All the photos were taken under a fluorescent microscope (mshot).

